# HapAdapter: A Transfer Learning Protocol for DNA Foundation Models Adapted to Diploid Personal Epigenomes

**DOI:** 10.64898/2026.11.25.769310

**Authors:** Joel Rozowsky, Amy Metrick, Jacqueline Wang, Andrei Onut, Jeb Cui, Tianxiao Li, Mark Gerstein

**Author notes:** These authors contributed equally to this paper (Joel Rozowsky, Amy Metrick).

## Abstract

The rapid rise of large language models (LLMs) has spurred the development of genomic language models (gLMs) that capitalize on the key features of natural language present in biological sequences. Moreover, DNA foundation models, a sub-class of gLMs, have demonstrated impressive accuracy in predicting a wide range of functional tracks from DNA sequence input. However, current DNA foundation models are exclusively trained on haploid reference genome sequences – despite personal and graph genomes becoming the new standard for genomic analyses. This is principally because the cost to train gLMs on a personal diploid genome for a single individual is not currently practical. To help reconcile this disconnect between the current available use-cases of DNA foundation models with the modern direction of genomic sequence analysis, we present HapAdapter, a transfer-learning protocol that builds a small Siamese network on top of any pre-trained foundation model to enable inference from full diploid personal genome context. To demonstrate the utility of this method, we train a set of HapAdapter heads to predict allele-specific effects across assays on two representative foundation model backbones: DNABERT-2 and AlphaGenome. We demonstrate that our lightweight adapter more than doubles DNABERT-2’s allele-specific discrimination and triples AlphaGenome’s (130% and 210% gain in AUROC above chance, respectively, averaged over nine assays).

## Introduction

In recent years, there has been a high level of enthusiasm for applications of frontier models in the field of genomics (J. Zhou et al., 2018; Avsec et al., 2026; Brixi et al., 2026). As existing research (Searls, 2002; Ji et al., 2021) shows that DNA sequence shares many fundamental characteristics with human language (e.g. context-dependent meanings and long-range semantic interactions), this presents an opportunity to utilize natural language processing techniques for genomics applications. This is particularly helpful as while protein translation is well understood, the behavior of regulatory and non-coding regions vary across contexts.

In parallel, there has also been greater emphasis on using personal genomes for the analysis of matched functional genomics data (J. Rozowsky et al., 2023). This has become manifest with the evolution in sequence technologies and the ability at relatively modest expense to fully assemble personal diploid genome sequences. This is also evidenced by the development of the human pangenome (Liao et al., 2023) made possible by graph genome approaches (Computational Pan-Genomics Consortium, 2018; Paten et al., 2017) to better represent personal genomes. In this paper we demonstrate how frontier genomic language models can be used for inference on diploid personal genomes. In particular, we develop a transfer learning protocol and apply it to the problem of predicting allele-specific activity.

Predicting allele-specific (AS) behavior is an ideal problem for the application of frontier models trained on personal epigenome data. Traditional approaches for accurately determining allele-specific behavior require deeply sequenced functional genomic data, such as RNA-seq or ChIP-seq. This functional data must be derived from biological material sourced from an individual that has also been fully genotyped to enable matched construction of a diploid personal genome for all chromosomes (Joel Rozowsky et al., 2011; J. Rozowsky et al., 2023). Mapping these reads to the personal genome enables the identification of reads that overlap heterozygous variants, where maternally and paternally derived haplotypes differ. Sites or regions exhibiting allele-specific activity can then be classified by determining loci that exhibit statistically significant differences (Joel Rozowsky et al., 2011) between the activity (such as transcription or transcription factor (TF) binding) on the two haplotypes overlapping heterozygous variants.

We present HapAdapter, a transfer learning protocol that adapts pre-trained genomic foundation models to enable accurate personal-genome based inference. We train an array of HapAdapter heads using data derived from the EN-TEx resource of matched personal epigenomes (J. Rozowsky et al., 2023) to accurately predict sites of allele-specific activity surrounding heterozygous SNVs (Zhang et al., 2021; Cleary and Seoighe, 2021). We demonstrate how this protocol can be implemented on two frontier model backbones: DNABERT-2 (Z. Zhou et al., 2024) and AlphaGenome (Avsec et al., 2026). We also show how the resulting HapAdapter heads on both frontier backbones generalize to unseen individuals at inference time. Furthermore, we demonstrate that this approach is effective for predicting different types of binding activity, such as transcription factor (TF) binding and histone modifications. In addition, we compare the performance of this protocol when the DNA sequence windows are extracted from the personal relative to the reference genome.

Using in silico saturation mutagenesis (Alipanahi et al., 2015; J.Zhou and Troyanskaya, 2015), we further investigate the DNA sequence features that the resulting models are most sensitive to in predicting AS behavior. We compare these features with motifs of known transcription factors (Ovek Baydar et al., 2026).

## Results

### The HapAdapter Protocol

HapAdapter (see Figure 1) is an efficient transfer learning protocol that enables any DNA foundation model trained on haploid reference genomes to be adapted for diploid personal genome inference. For a set of genomic windows, the protocol extracts final-layer embeddings from the chosen foundation model through one forward pass for each haplotype and then trains the HapAdapter head on the resulting haplotype embedding pairs. The HapAdapter head architecture follows the “siamese” network formulation first proposed by Bromley et al., 1993.

**Figure 1.**
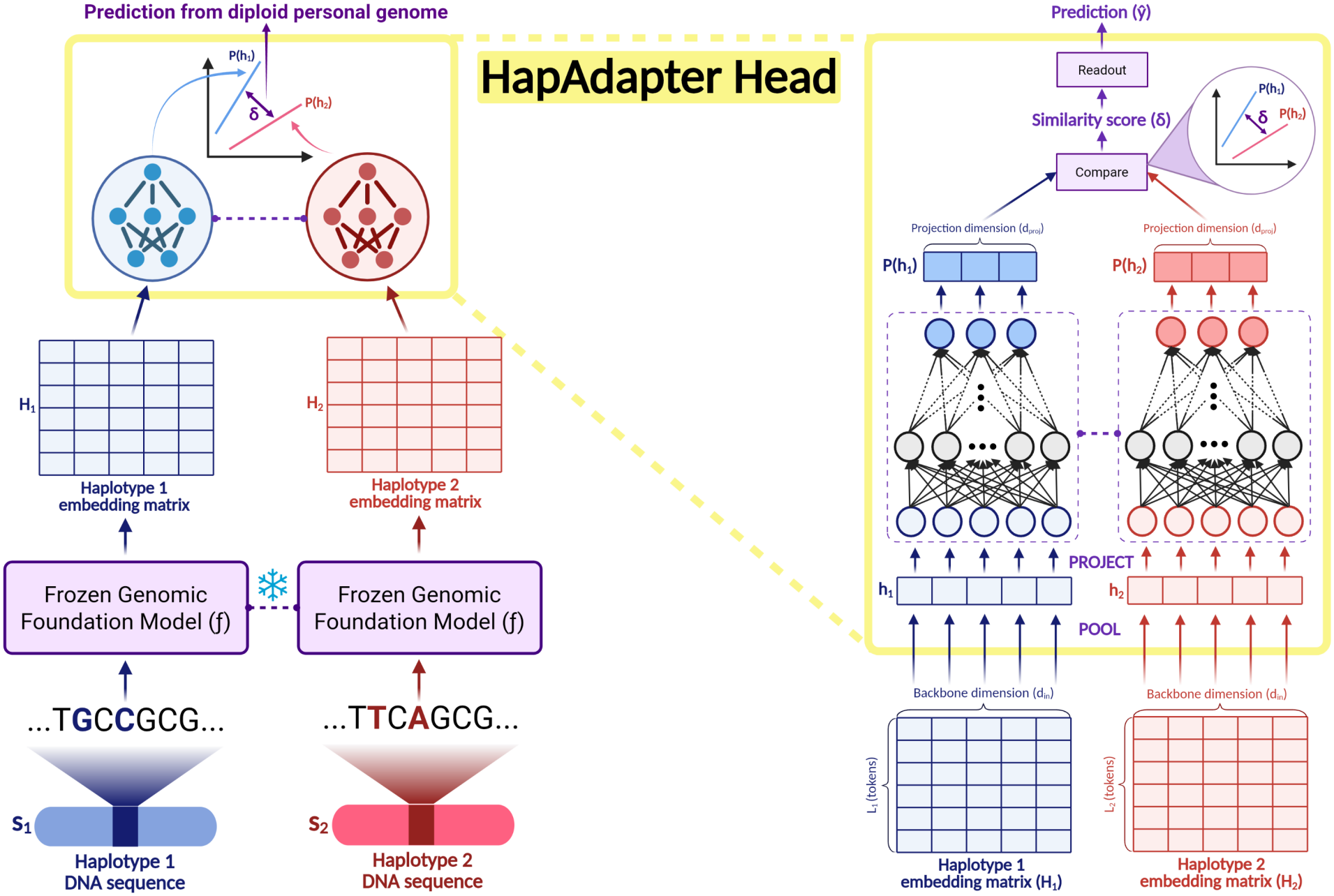
Schematic of the HapAdapter Protocol. A schematic of the HapAdapter transfer learning protocol, which trains a “siamese” network head to predict a functional track (e.g. the presence of allele-specific activity at a variant) from the extracted final-layer (haplotype 1, haplotype 2) embeddings provided by a frozen genomic foundation model. The right hand side zooms in on the details of the HapAdapter head, including pooling of embeddings and learned projection into lower-dimensional space that is then used by the head for discrimination via a “similarity score”. Created in BioRender. Metrick, A. (2026) https://BioRender.com/gy9r80v.

In particular, let *s*_1_ and *s*_2_ denote the two input haplotype sequences for a given locus, which are then tokenized into *L*_1_ and *L*_2_ tokens respectively (note that *L*_1_ is not necessarily equal to *L*_2_ in all tokenization schemes). Let *f* be the frozen foundation model of choice with intrinsic embedding dimension *d*_in_. Then after one forward pass through the foundation model, we can extract the two matrices of final-layer learned token embeddings associated with each of the two haplotypes (call them *H*_1_ and *H*_2_ respectively):

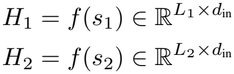

The HapAdapter head takes in these matrices and first pools across tokens within each of the *d*_in_ feature dimensions. Note that this can be done with any reasonable pooling function (e.g. max pooling, min pooling, mean pooling, etc), but we have chosen center-mean pooling (i.e. averaging the center few tokens within each feature dimension) in our implementation as we center the variants in the input windows. After pooling, we get the flattened haplotype embedding vectors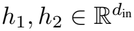:

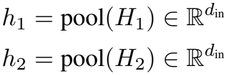

These pooled embedding vectors then enter a shared Multi-Layer-Perceptron (MLP) network that is trained to project them into a lower-dimensional space against a specific objective function (see Application: Predicting Allele-Specific Effects for an example objective). The resulting lower-dimensional embedding vectors are then compared in this lower-dimensional space (e.g. using Euclidean distance or cosine similarity) and the resulting similarity score is read out through the objective function the projection MLP was trained on to obtain a prediction.

For backbone foundation models that are pre-trained on sequence reconstruction alone (i.e. no pre-training to predict functional tracks), we recommend first fine-tuning the backbone on a related functional task using reference genome sequences (see Figure S1). We employed this method for the DNABERT-2 model (see Methods).

### Application: Predicting Allele-Specific Effects

#### HapAdapter AS Objective Function

By definition, a locus that exhibits allele-specific activity sees a statistically significant difference in behavior at each of the two parental haplotypes. We would thus expect that the embeddings for the two haplotype sequences at an AS locus should be more different from one another than the two haplotype embeddings at a non-AS (balanced) locus. We designed the objective function for predicting the presence of AS activity to reflect this intuition.

Let *P* denote the projection of pooled haplotype embedding vectors into lower-dimensional space that we aim to train in the HapAdapter head. We would like *∥P* (*s*_1_) *™P* (*s*_2_) *∥* to be large when *s*_1_ and *s*_2_ are an AS haplotype pair and small otherwise so that it is easy for the model to classify AS behavior just by observing the distance between the projected haplotype embeddings.

We define the *dissimilarity* score between the two haplotype embeddings to be the Euclidean distance *δ* between their images under the projection *P* :

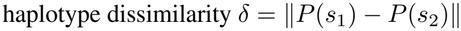

Two scalar parameters, *a* and *b*, are learned jointly with *P* in the HapAdapter head and exist to map *δ* to a logit:

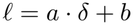

Then the probability of AS behavior that the model assigns to the input haplotype sequence pair (*s*_1_, *s*_2_) is:

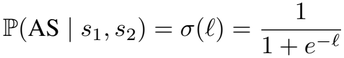

The model then predicts AS behavior when it assigns a probability of greater than 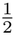 to that sequence exhibiting AS effects. In particular, when 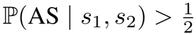 (which coincides with *ℓ >* 0 since 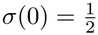). Equivalently, the model classifies the haplotype pair as allele-specific when 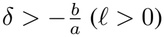 and not allele-specific when 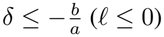:

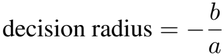

So the model prediction can be written as:

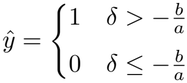

where 1 means AS and 0 means non-AS. The objective function for training the HapAdapter head is then the negative of Binary Cross-Entropy (BCE) loss:

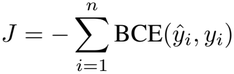

where there are *n* training sequences, the *i*-th of which with true label *y*_*i*_ and predicted label *ŷ*. The code optionally supports weighting the loss by experimental read-depth of the input sequences. This is done in the published versions of the models to prevent throwing away data built from experiments with modest read-counts while still ensuring that the model’s gradients don’t move too drastically on less well-powered examples. Concretely,

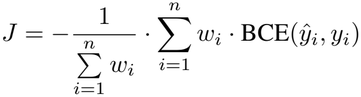

where *w*_*i*_ is the precision-weight given to example *i*, which is inversely proportionally to the read-depth at that locus:

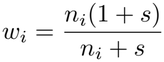

Here, *n*_*i*_ is the number of reads for locus *i* and *s* is the saturation constant ensuring that the precision-weight saturates once a locus reaches a sufficiently large read-depth.

In essence, this objective “pulls apart” the two haplotype embeddings for loci that exhibit AS effects and “pulls together” the two haplotype embeddings for loci that do not exhibit AS effects. We can see this desired separation emerge by comparing the distances between the two haplotype embeddings in AS examples vs. non AS examples from frontier model backbones to their corresponding trained HapAdapter heads (see Figure 2).

**Figure 2.**
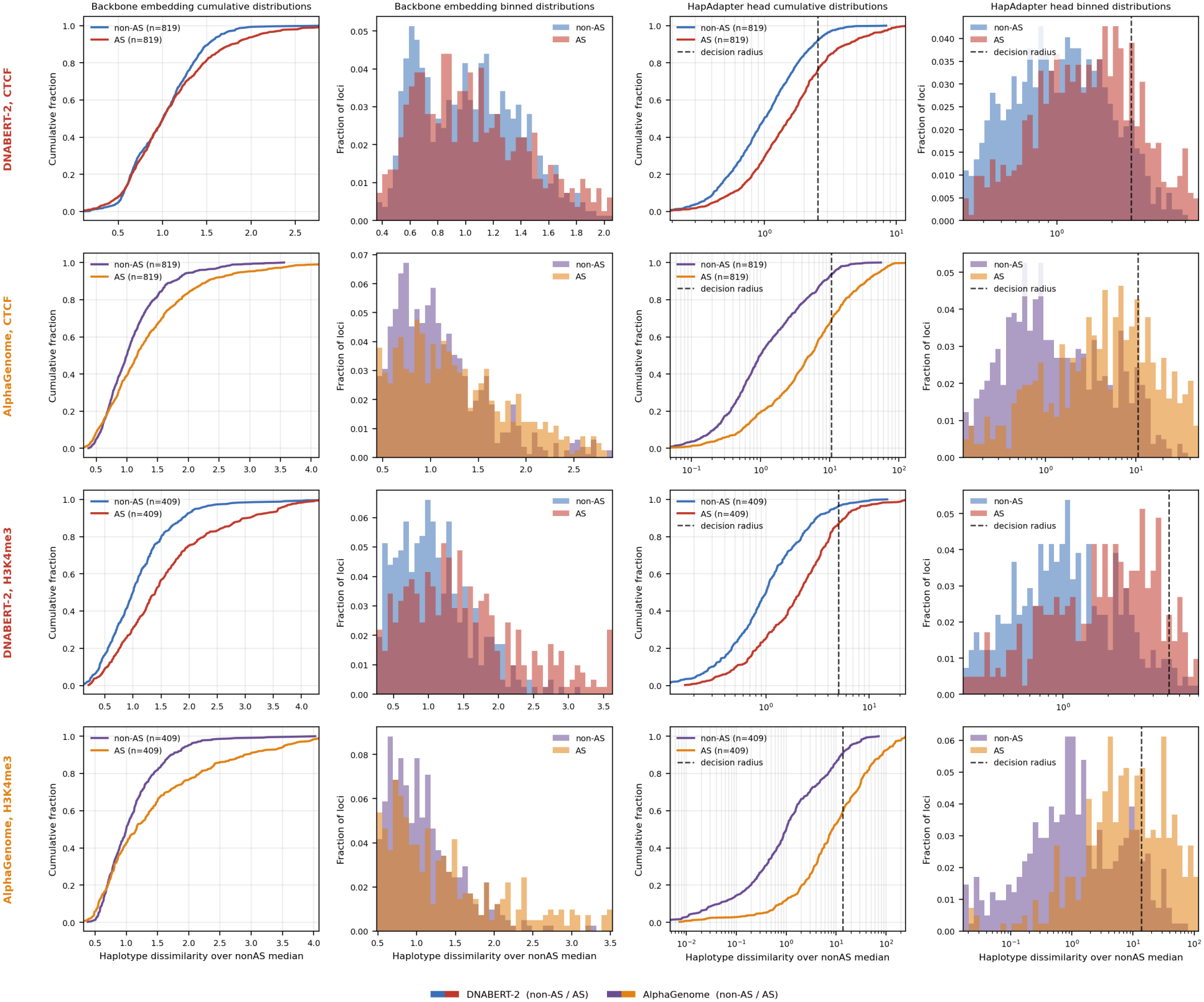
Comparison of the HapAdapter protocol for two foundation models. Visualization of the HapAdapter head in discriminating AS from non-AS haplotype pairs for two representative assays (CTCF and H3K4me3 ChIP-seq from the EN-TEx resource), using backbones from two foundation models (DNABERT-2 and AlphaGenome). Rows are (backbone, assay) pairs for individual 2. The leftmost two panels show the haplotype dissimilarity read from the frozen foundation-model backbone alone, as a cumulative distribution and then as a binned distribution. The right two panels show the same quantity after the trained HapAdapter head on a logarithmic axis, where the dashed line marks the head’s decision radius. Dissimilarity is expressed relative to each row’s non-AS median and bars give the fraction of each class’s loci per bin. Loci are drawn from each assay’s AlphaGenome balanced positive/negative test split. The head panels are clipped for display: each axis spans the 3rd to 97th percentile of that row’s dissimilarity.

#### Two Frontier Model Backbones: DNABERT-2 and AlphaGenome

DNABERT-2 is pre-trained with the masked language modeling objective (MLM) on DNA sequences, and sees no functional labels during pre-training (Z. Zhou et al., 2024). The MLM objective is intended to enable the model to learn general DNA “grammatical” patterns. However, it is not necessarily going to focus on learning types of *binding* or other specific functionally-linked grammars in this unsupervised setting. Thus, before fitting the HapAdapter head on the DNABERT-2 backbone, we fine-tuned DNABERT-2 to predict binding affinity for each assay we planned to fit the AS HapAdapter head on. This form of intermediate-task transfer (Phang et al., 2018; Pruksachatkun et al., 2020) allows the model to intuitively bridge the gap from sequence-level understanding to making a functional prediction based on a diploid personal genome by first modifying its learned representation of DNA sequences to better capture the relevant binding grammar on reference genome sequences. In its own right, we see that DNABERT-2 fine-tuned for regression with precision-weighted MSE loss performs well for predicting binding strength across assays and individuals (see Figure S2).

On the corresponding fine-tuned DNABERT-2 backbones, HapAdapter heads were built and tested for 11 assays across the four EN-TEx individuals for both “reference” and “personal” arms. The reference-sequence heads saw (ref, alt) input pairs, where ref was the reference genome sequence surrounding the given variant and alt was the reference sequence with the SNV nucleotide switched to the alternate allele. The personal-sequence heads saw the full (hap1, hap2) diploid personal genome sequences. The personal sequence heads performed an average increase of 0.041 AUROC over the reference sequence heads across assays and individuals (see Figure 3A). The model performance only saw a drop of about -0.0025 AUROC across assays (see Figure 3B) when scored on the non-training individuals, highlighting impressive generalization to unseen individuals despite being fit to one individual’s personal genome.

**Figure 3.**
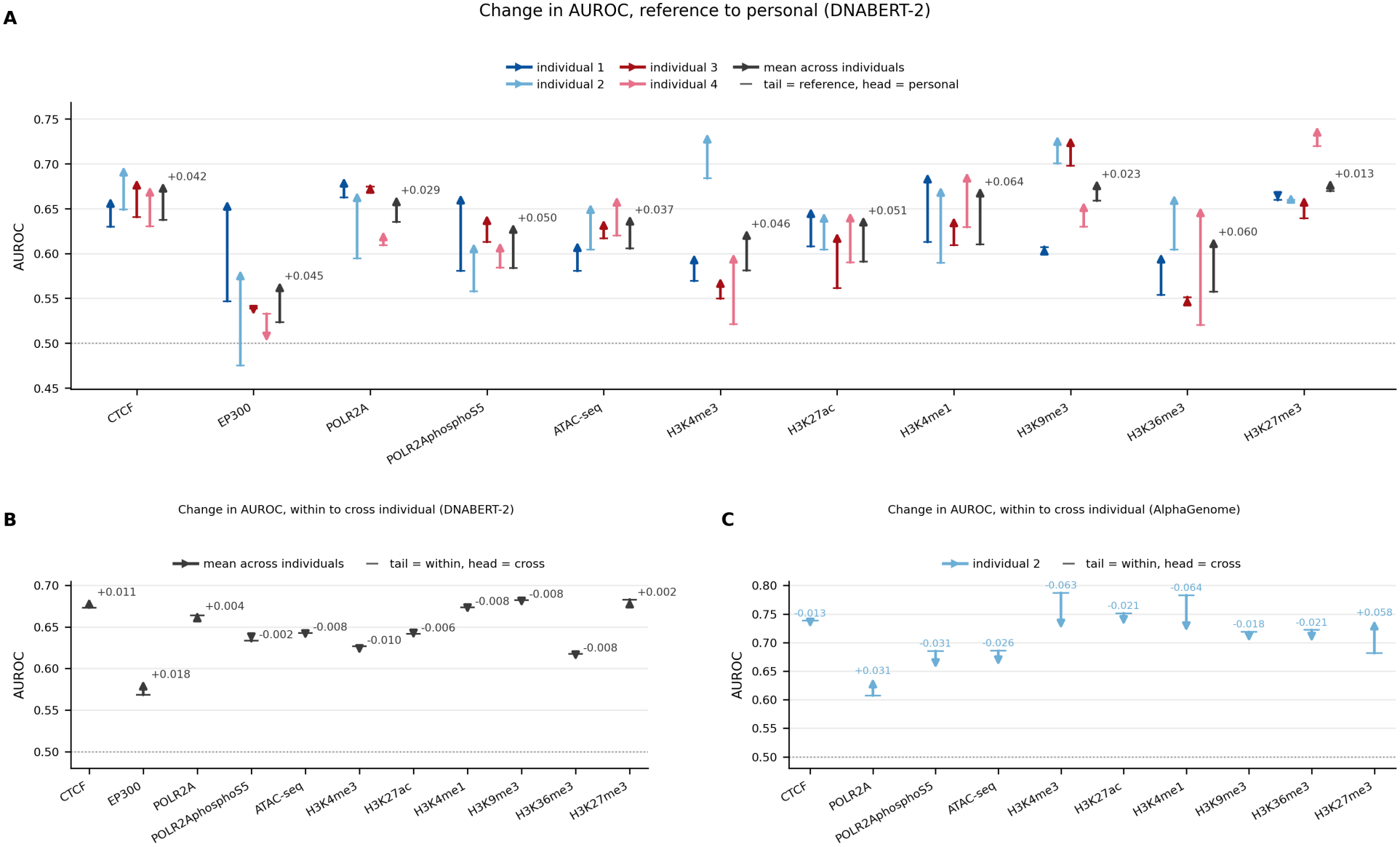
HapAdapter performance comparisons. **(A)** Difference in HapAdapter performance (AUROC) using the DNABERT-2 backbone across eleven EN-TEx assays and four individuals, between using reference genome sequences (with the variant replaced in the paired sequence) and the personal diploid sequence flanking the hetSNV of interest. Each arrow runs from the reference value (tail) to the personal value (head). Color denotes individual and the black arrow is the mean across individuals. **(B)** A head trained on one individual transfers to the remaining individuals with minimal loss: arrows run from the within-individual AUROC (tail) to the mean across the three held-out individuals (head), for the DNABERT-2 backbone. **(C)** The same comparison for AlphaGenome, for HapAdapter heads trained on individual 2 and transfer performance averaged over the other three EN-TEx individuals.

Unlike DNABERT-2, the pre-training stage of AlphaGenome includes functional task supervision (Avsec et al., 2026). Thus, we skipped fine-tuning the AlphaGenome backbone and proceeded straight to fitting the HapAdapter head on 10 assays for individual 2 only (excluding the EP300 assay that came up underpowered once positives and negatives were balanced). The AlphaGenome-based HapAdapter heads also demonstrated impressive generalization across individuals, with an average performance drop of -0.0168 AUROC across assays when scoring models trained on individual 2 on the other three donors (see Figure 3C).

#### The HapAdapter Performance Boost

To examine whether the HapAdapter head improves performance on predicting allele-specific activity compared to the pre-trained foundation models when the diploid personal genomes are provided, we evaluate and compare several reasonable ways of using the foundation models in isolation for predicting the presence of AS effects.

First, we evaluate the performance of each foundation model when used to classify-on-distance from the two haplotype final-layer embeddings as done in the objective function of the HapAdapter head. For DNABERT-2, we compute both the performance for the pre-trained backbone (that has only learned sequence-level patterns by construction) and the backbone after being fine-tuned to predict the appropriate binding affinity. Since AlphaGenome is already pre-trained with functional task supervision, there is no analogous “sequence only” evaluation to run. These evaluations are primarily done to directly compare the improvement brought about by the HapAdapter head trained with the objective focused directly on these embedding space distances.

In addition, we evaluate the more natural zero-shot baseline, computed by differencing the actual predicted binding tracks for each haplotype from the fine-tuned DNABERT-2 backbone and the AlphaGenome backbone (rather than differencing their final-layer embeddings).

In both cases, we see the HapAdapter head substantially improves performance in classifying AS activity across assays (see Figure 4). Compared to the corresponding zero-shot baselines, the HapAdapter heads on average more than double DNABERT-2’s allele-specific discrimination and triple AlphaGenome’s.

**Figure 4.**
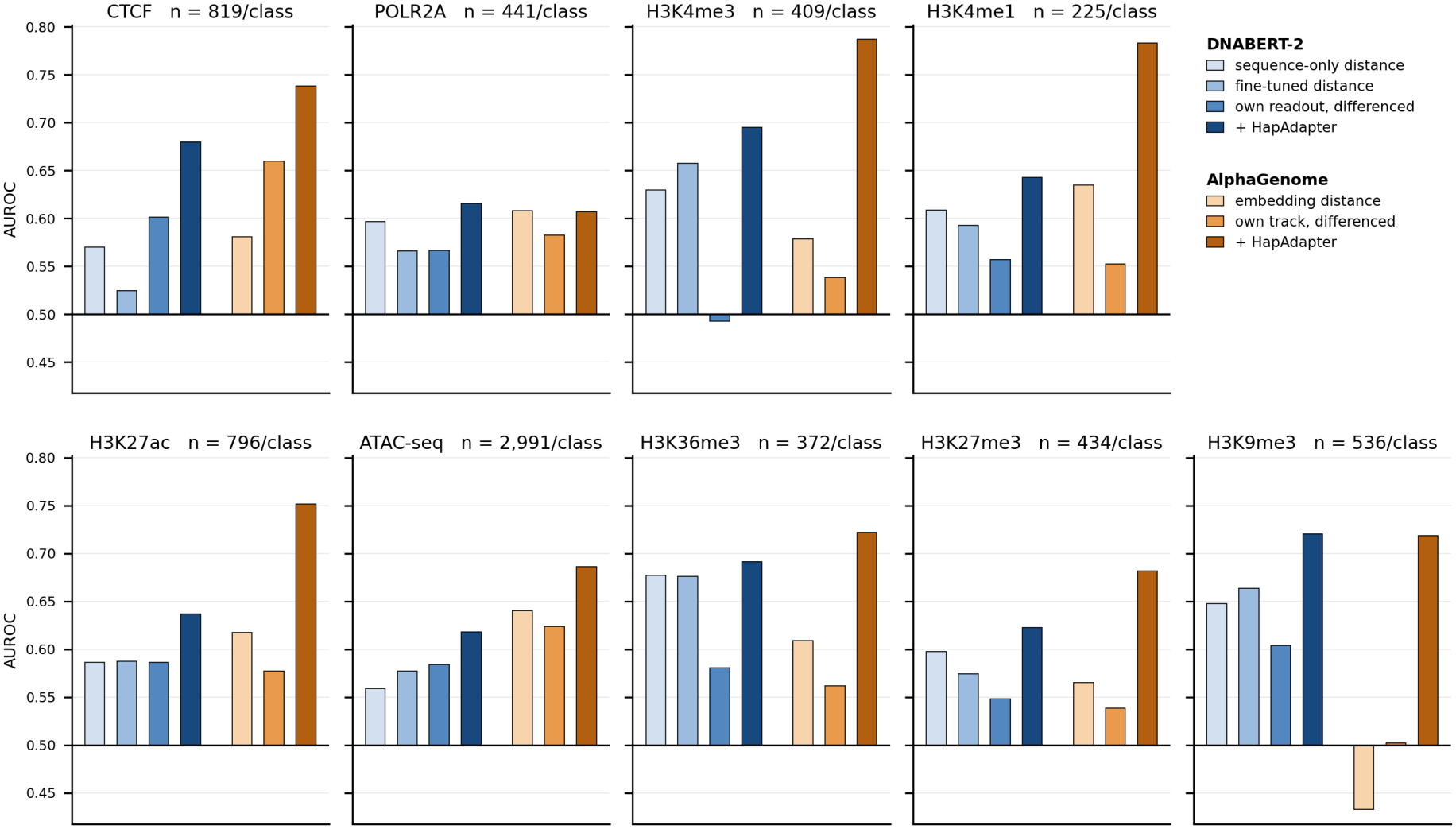
Comparison of backbone and HapAdapter head performance across assays and frontier models. AUROC for discriminating AS from non-AS haplotype pairs across nine EN-TEx assays for individual 2, using the DNABERT-2 and AlphaGenome backbones. Within each panel, bars to the left are DNABERT-2 and to the right AlphaGenome. Within each family the bars are: the sequence-only embedding distance (DNABERT-2 only), the functionally supervised embedding distance, the model’s own native allelic-difference readout, and the trained HapAdapter head. The HapAdapter head outperforms both the embedding distances and as well as each backbone’s own native variant-effect readout across all nine assays. The trained AlphaGenome HapAdapter heads outperform the trained DNABERT-2 HapAdapter heads on eight out of the nine assays.

#### ISM & Motif Enrichment

To elucidate which DNA sequence features most significantly impact model prediction, we perform in silico saturation mutagenesis (ISM) experiments (Alipanahi et al., 2015; J. Zhou and Troyanskaya, 2015) on the DNABERT-2 HapAdapter heads, measuring how much each model changes its prediction in response to every single nucleotide substitution across the test sequences (see Methods). We find that the resulting ISM saliency profile plots peak at the hetSNV of interest, with true positive classified sequences consistently showing the highest central peak across assays and individuals (see Figure S3).

We further examined the ISM profile of CTCF, a highly-conserved transcription factor with a well-documented binding motif and cofactor complex. As seen across assays and individuals, the ISM profile has the highest central peak for the true positive sequences, followed by false positive, false negative, and finally true negative sequences (see Figure 5A). Strikingly, the ISM contribution logo averaged across allele-specific sequences surrounding the central hetSNV matches up closely with the true CTCF binding motif (see Figure 5C).

**Figure 5.**
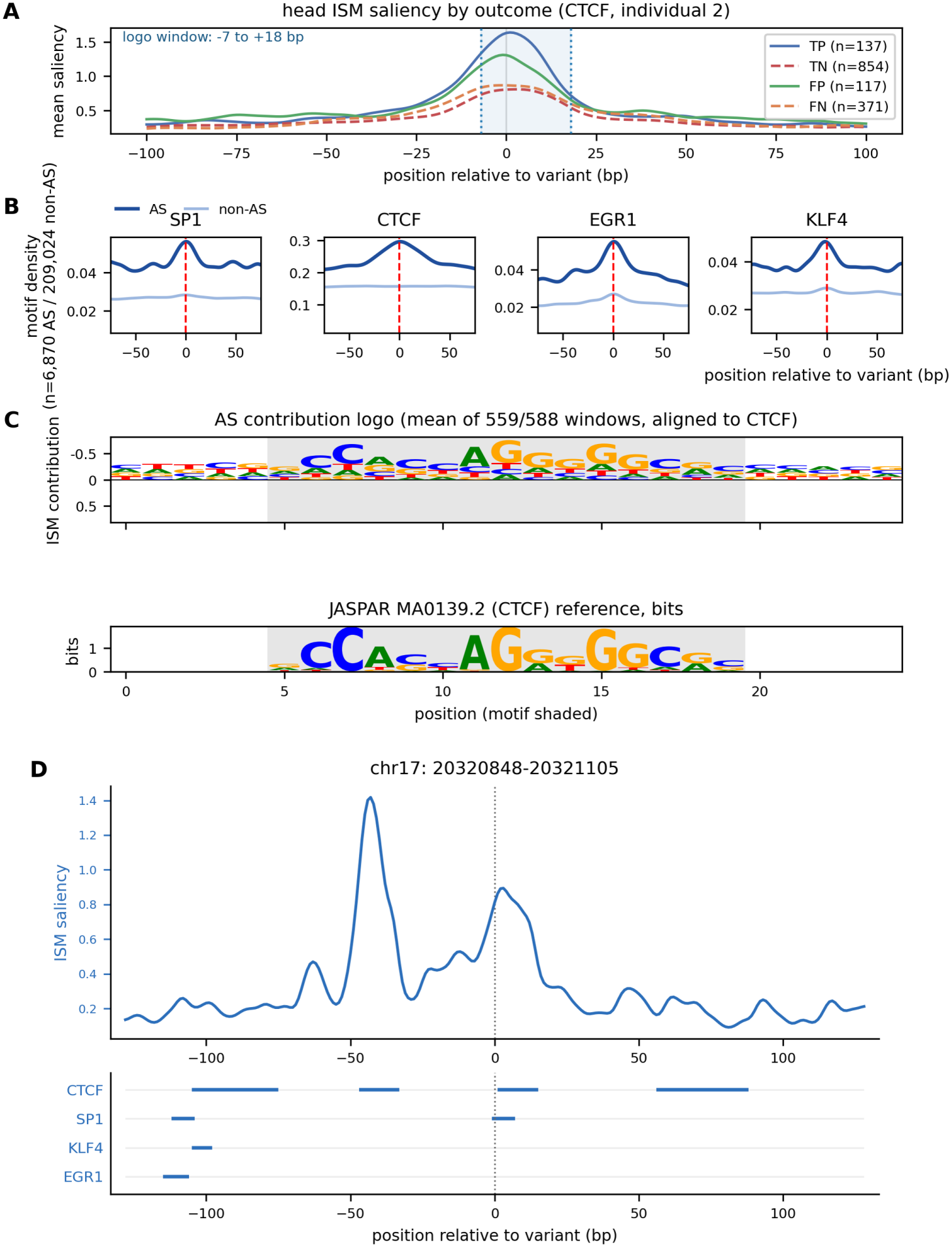
In silico saturation mutagenesis of the DNABERT-2 HapAdapter head and analysis of associate TF motif enrichment. **(A)** ISM saliency of the HapAdapter head on the DNABERT-2 backbone for CTCF, individual 2, averaged within each classification outcome. The head is most responsive to mutations at the center of the window (where the variant is located), and separates true positives and false positives from false negatives and true negatives. The shaded band marks the span contributing to the contribution logo in (C). **(B)** Density of CTCF and known cofactor motifs (SP1, EGR1, KLF4) as a function of position relative to the variant (marked by a dashed line), for AS and non-AS loci separately. **(C)** Mean ISM contribution across AS windows, aligned to the CTCF motif, with the JASPAR MA0139.2 reference shown underneath (Ovek Baydar et al., 2026). The vertical axis is inverted so that *negative* ISM contribution points upward: mutating a functional base reduces the predicted quantity, so the bases the model relies on carry negative contribution, and plotting them upward makes the recovered motif readable. **(D)** A representative locus classified as allele-specific for CTCF binding (chr17:20,320,848–20,321,105), showing the ISM saliency profile and the positions of CTCF and cofactor motif hits in the same sequence.

Furthermore, we examined single windows across the CTCF test set and discovered that CTCF as well as CTCF cofactor binding motifs matched up with peaks in the saliency profiles (one example is shown in Figure 5D). Examining these cofactors further, we notice elevated motif density on average at the central hetSNV location within the training and testing datasets (see Figure 5B).

### Documentation and Distribution via GitHub Repository and Hugging Face

All data used for model training, evaluation, and testing comes from the EN-TEx resource, which is available on the EN-TEx portal located at https://entex.encodeproject.org. The HapAdapter source code is available on GitHub with documentation detailing usage at https://github.com/gersteinlab/HapAdapter. The trained models are available on Hugging Face as a collection of eleven repositories, one per assay, at https://huggingface.co/collections/ametrick-system/hapadapter.

## Discussion

In this paper, we outline the HapAdapter transfer learning protocol for adapting frontier genomic language models to perform personal variant-aware inference on phased diploid genome sequences. The protocol trains a siamese network (Bromley et al., 1993) on extracted final-layer haplotype embeddings from frozen pre-trained DNA foundation models. Thus, this technique can be used to adapt any foundation model with accessible final-layer embeddings to make individualized predictions when a diploid personal genome assembly is available.

We demonstrate the efficacy of this method for predicting allele-specific activity on two foundation model backbones: DNABERT-2 and AlphaGenome. We show that this approach achieves ever greater improved accuracy when using personal diploid genome sequences rather than the reference-alternate sequence pairs. We also show that despite being trained exclusively on one individual’s personal genome sequence, the HapAdapter heads transfer well between individuals at inference time, suggesting that they learn how to compare haplotype pairs within individuals without overfitting to the training individual. Furthermore, we demonstrate that the trained HapAdapter heads are most sensitive to perturbations in the sequence features that lie within relevant cofactor motif regions via an in silico saturation mutagenesis approach.

There are several limitations to our approach that are worth highlighting to inform future work. Our instantiation of the HapAdapter head for predicting AS effects is restricted to a binary readout by construction, as the predicted quantity is based on the distance between the two haplotype embeddings, which is non-negative and symmetric. In particular, it cannot express a signed effect and predict which haplotype is favored in the allelic imbalance. However, we note that our general framework can be extended to continuous or multi-bin targets if the objective function is altered. We present this approach as our proof-of-concept example due to the nice geometric intuition that comes with it. Furthermore, though the EN-TEx individuals are selected to be diverse, our evaluation does rest on four individuals from a single resource. We hypothesize that our method would still generalize to other individuals from distinct datasets, but we have not yet tested this hypothesis.

We believe these limitations do not detract from the methods contribution of HapAdapter in bridging the gap from DNA foundation models to personal genomic analysis. We plan to continue investigating ways to apply the HapAdapter protocol to predict a range of personalized functional tasks.

## Methods

### Using data from the EN-TEx personal epigenome resource

Matched personal genomes and functional genomics data used in this paper for training and testing the suite of HapAdapter heads to predict allele-specific effects came from the EN-TEx resource of personal epigenomes (J. Rozowsky et al., 2023). The EN-TEx resource includes the fully assembled diploid personal genome sequences for 4 individuals (2 male and 2 female) along with the matched functional genomic assays for approximately 30 different tissues and 10 assays (e.g. RNA-seq, ChIP-seq, ATAC-seq, Hi-C, and so on). These functional assays were mapped and summarized in bigWig files quantifying the amount of functional activity across the genome in these individuals from both the human reference genome sequence (hg38) and the matching personal diploid genome sequences.

In addition, the mapped reads for each assay were processed using the AlleleSeq pipeline (Joel Rozowsky et al., 2011) to determine sites of allele-specific activity. For each functionally active heterozygous SNV, this pipeline classifies its allele-specific status and outputs the read counts corresponding to each haplotype along with the p-value evaluating it’s allelic classification. This output was used for training and testing of the HapAdapter AS heads as detailed below.

### Extraction and labeling of DNA sequence windows for training the HapAdapter AS classifiers

Candidate sites for model training and testing were the heterozygous SNVs from the `hetSNVs_default_AS.tsv` build on the EN-TEx portal (http://entex.encodeproject.org/main.html). For each donor and assay, allelic read counts were summed across tissues and a locus was labeled allele-specific if any tissue called significant imbalance.

All the DNABERT-2 HapAdapter heads were trained on 257 bp windows, each centered on a hetSNV and labeled with a 1 if the hetSNV exhibited AS effects for the given assay and individual in any catalogued tissue and 0 otherwise. Four sequences were extracted for each hetSNV locus per individual: (1) reference genome sequence window (hg38),

(2) reference genome sequence window with only the hetSNV swapped for the alternate allele (referred to as the “alt” sequence), (3) paternal haplotype sequence window, and (4) maternal haplotype sequence window. The haplotype sequences were extracted by lifting the hetSNV coordinate through that individual’s chain file, with loci that failed to lift, fell in an assembly gap, or contained an ambiguous code (e.g. “N”) discarded. A HapAdapter head was trained for each assay on both the (ref, alt) sequence pairs and the personal genome haplotype sequence pairs and the performance of the two heads were compared (see Figure 3).

The HapAdapter heads were trained, selected, and evaluated using an approximate 8:1:1 split of the full dataset for a given assay and individual. The training split made up the *∼*80%, while the development set used to select the best model checkpoint made up *∼*10%. Finally, the other *∼*10% made up the testing set, which was composed of entirely held-out chromosomes and never seen during the model training process.

These train/dev/test splits for each assay were assigned per locus and remained consistent for hetSNVs that appeared in the dataset for more than one donor to ensure leak-free cross-donor evaluation. This was done by keying on the SHA1 (National Institute of Standards and Technology, 2015) hash of chromosome and position per locus. For each assay, whole chromosomes were held out for testing, chosen deterministically to cover about 10% of the positive sequences, excluding the sex chromosomes, chromosome 6, and the acrocentric chromosomes, as well as the two chromosomes with the most positive loci in the given dataset. Remaining chromosomes were divided tiled into 100 kb bins, and each bin was assigned entirely to the train or dev set in an 8:1 ratio via SHA1 hashing, with sequences within 128 bp of a bin edge discarded to avoid any sequence overlap between the train and dev sets.

AlphaGenome accepts only a fixed set of longer input lengths, so its corresponding HapAdapter heads used 16,384 bp windows re-extracted at the same loci: 8,192 bp left of the hetSNV and 8,191 bp right, placing the variant one base right of center. Labels and train/dev/test assignment were inherited unchanged from the 257 bp build for DNABERT-2, so both backbones are compared on the same variants. Because a 16,384 bp window reaches well beyond the 128 bp bin margin, loci whose widened window overlapped a window in another split were dropped.

The two backbones were not trained on identically composed sets due to the computational power required to extract AlphaGenome embeddings for each sequence. The DNABERT-2 heads were fitted on the full build at each split’s natural class prevalence, whereas the AlphaGenome windows were re-extracted as a 1:1 subsample in which each positive was paired with a negative drawn from within the positives’ read-depth deciles, applied to all three splits. Any comparison between backbones (Fig. 4) was therefore made after restricting both arms to their shared locus set.

### Extraction of foundation model embeddings for input into the HapAdapter heads

Each haplotype window was passed through each foundation model once and reduced to a single pooled embedding vector for training the associated HapAdapter heads. For DNABERT-2, the windows were tokenized using the model’s own byte-pair encoding tokenizer. Center-mean pooling was then applied by averaging the final hidden states of the five tokens spanning the middle of the window, yielding a 768-dimensional pooled embedding vector per haplotype. For AlphaGenome, the single-base-resolution trunk was used and the resulting embeddings were also center-mean pooled over the 5 central tokens (which are individual nucleotides here), yielding a 1536-dimensional pooled embedding vector per haplotype. Pooled vectors then enter the head as the pair (*h*_1_, *h*_2_) with no further normalization.

### Functional fine-tuning of the DNABERT-2 backbone

Some genomic foundation models are not pre-trained to predict any functional genomics tracks. Rather, they are exclusively pre-trained to predict missing segments of DNA sequences themselves, thus prioritizing the model’s understanding of basic sequence semantics without linking these patterns to any functional readout. This is the case for DNABERT-2 (Z. Zhou et al., 2024), which pre-trains on the Masked Language Modeling (MLM) objective introduced in the original formulation of the BERT (Bidirectional Encoder Representations from Transformers) architecture (Devlin et al., 2019). We recommend fine-tuning these models on a relevant functional task before fitting the HapAdapter head to the backbone.

Our method used on DNABERT-2 works for any pre-trained encoder network that can be instantiated through the HuggingFace `AutoModel` interface from a single `–model_name_or_path` argument. The only backbone-dependent quantity is the intrinsic model dimension *d*_in_, which is read from the loaded configuration (*d*_in_ = 768 for DNABERT-2).

In our chosen application of predicting allele-specific binding using the HapAdapter protocol, we fine-tuned the DNABERT-2 backbone to predict the relevant binding track before fitting the HapAdapter head. That is, for each of the 11 assays we trained HapAdapter heads for on DNABERT-2, the DNABERT-2 backbone was fine-tuned to create an assay-specific trunk.

Trunk fine-tuning windows were anchored on binding peaks rather than on variants, and were built once per assay from the narrowPeak calls and fold-change-over-control bigWig signal tracks of every tissue catalogued for that assay. Peaks from all tissues were pooled and clustered by single-linkage chaining of their summits (summits within 100 bp of one another were merged into one consensus locus, represented by the summit of its highest-scoring member). Each consensus peak locus was paired with one background locus, sampled at random in proportion to each chromosome’s peak count and rejected within 1000 bp of any peak cluster.

Labels were derived from the fold-change-over-control bigWigs as the mean signal over the *±*32 bp centered on the anchor and averaged over tissues. The trunk was then fine-tuned as a regression over log(1 + FC) on 257 bp reference windows centered on the same anchors, using the reference genome only. The train/dev/test splits were determined via the same method as the AS head splits.

### Selection of hyperparameters

Most hyperparameters were inherited from established pipelines. Head batch size (32), warmup (50 steps), weight decay (0.01), and the trunk learning rate (3 *·* 10^*™*5^) follow the fine-tuning configuration reported for DNABERT-2 (Z. Zhou et al., 2024), which was grid-searched in that work and held fixed across all of its datasets. The trunk was fine-tuned at an effective batch of 128 (16 per device with 8 accumulation steps), and the HapAdapter heads were trained at a learning rate of 1*·*10^*−*3^. The HapAdapter head was chosen to be 2 layers deep to maximize computational efficiency while still allowing the model to capture non-linear features. The hidden and projection dimensions were both set to be 128. GELU activation, dropout 0.1, and depth-weighting constant *s* = 20 were further selected. Heads were trained for 3 epochs on DNABERT-2 embeddings and 25 on AlphaGenome embeddings to adjust for the difference in backbone and window sizes, with class-balanced batches and seed 20. The DNABERT-2 trunk was fine-tuned for 3 epochs with seed 42.

### Evaluation of model performance

Heads were selected on the dev split by AUROC, evaluated every 250 steps during training, and the best-performing checkpoint was used for scoring on the test set. All figures come from scoring on the held-out test chromosomes. Because class prevalence varies by assay, AUROC is also reported after subsampling the majority class to the size of the minority class.

### In silico saturation mutagenesis (ISM)

To shed light on the impactful sequence features on the HapAdapter head predictions, we performed ISM on the paired haplotype windows. This was done by substituting every alternate base at every position of the first haplotype window, holding the second window fixed as its partner, and measuring the change in the head’s prediction:

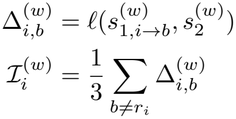

where *ℓ*(*s* , *s*) = *a ∥P* (pool(*s*))−*P* (pool(*s*)) *∥*+ *b* is the head’s logit for paired windows *s* , *s* , *w* is the window-pair index within the ISM testing set, 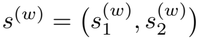 is the haplotype window pair for *w, b* is substituted base and *r*_*i*_ is the reference nucleotide at position *i* in 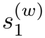. Thus, 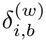 is the change in the logit when position *i* is substituted to base *b* in the haplotype 1 sequence and 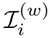 is the ISM attribution at position *i*, computed by taking the mean over the three possible substitutions. 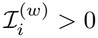 marks positions where substituting the reference base increases the predicted allelic contrast.

### Detection of motif enrichment

Position frequency matrices were taken from JASPAR (Ovek Baydar et al., 2026) and scanned against each window with FIMO (Grant et al., 2011) on both strands at a p-value threshold of 10^*−*3^ (deliberately looser than the typical 10^*−*4^ threshold used for genome-wide scans due to the short length of the windows). Motif density at a position is the fraction of windows in a group marked there, Gaussian smoothed for display. Densities are plotted as raw fractions on independent per-factor axes, since factors differ by an order of magnitude in how often they occur.

