## Supplemental Figures for "HapAdapter: A Transfer Learning Protocol for DNA Foundation Models Adapted to Diploid Personal Epigenomes"

### HapAdapter Supplement

### Supplemental Figures

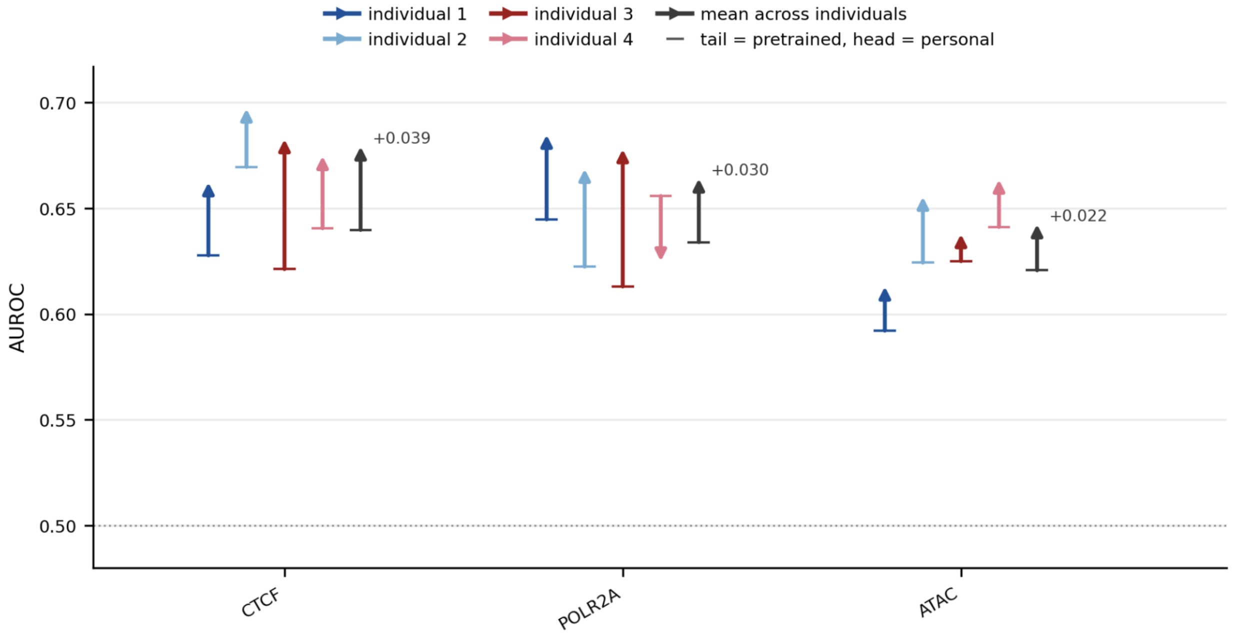


Figure S1. **Effect of the functional fine-tuning before fitting the HapAdapter head on DNABERT-2.** Before making the decision to run functional fine-tuning across assays and individuals for the DNABERT-2 models, we first compared the performance of the HapAdapter heads fit on the raw pre-trained backbones with the performance of the HapAdapter heads fit on the fine-tuned backbones for three representative assays (CTCF, POLR2A, ATAC-seq). We found an average increase in performance of about in 0.03 AUROC score, so we adopted the fine-tune first approach across assays and individuals.

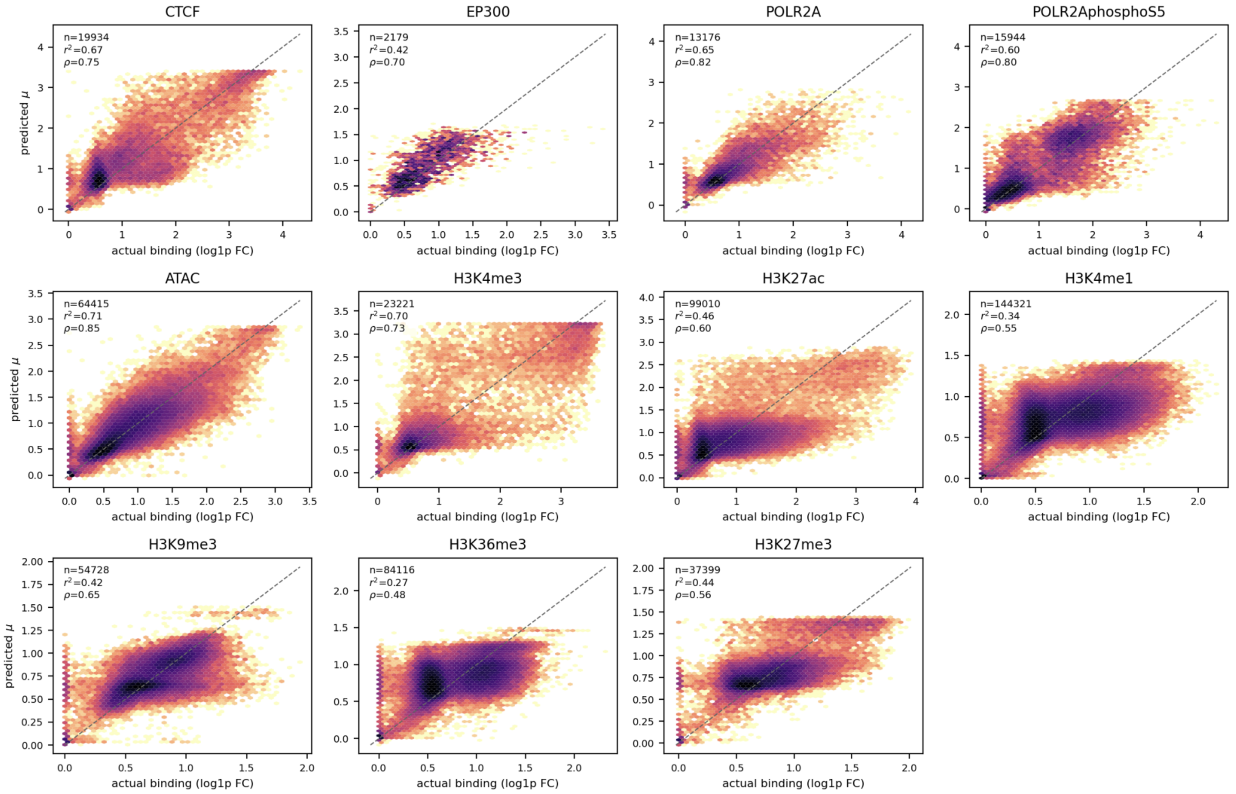


Figure S2. **Predicted vs. actual binding signal from DNABERT-2 backbone fine-tuning across assays for individual 2.** The fine-tuned intermediate model produced in the DNABERT-2 HapAdapter protocol pipeline performs reasonably well on predicting binding strength across assays and individuals, displayed here just for individual 2 as we observe the same behavior for the other three individuals.

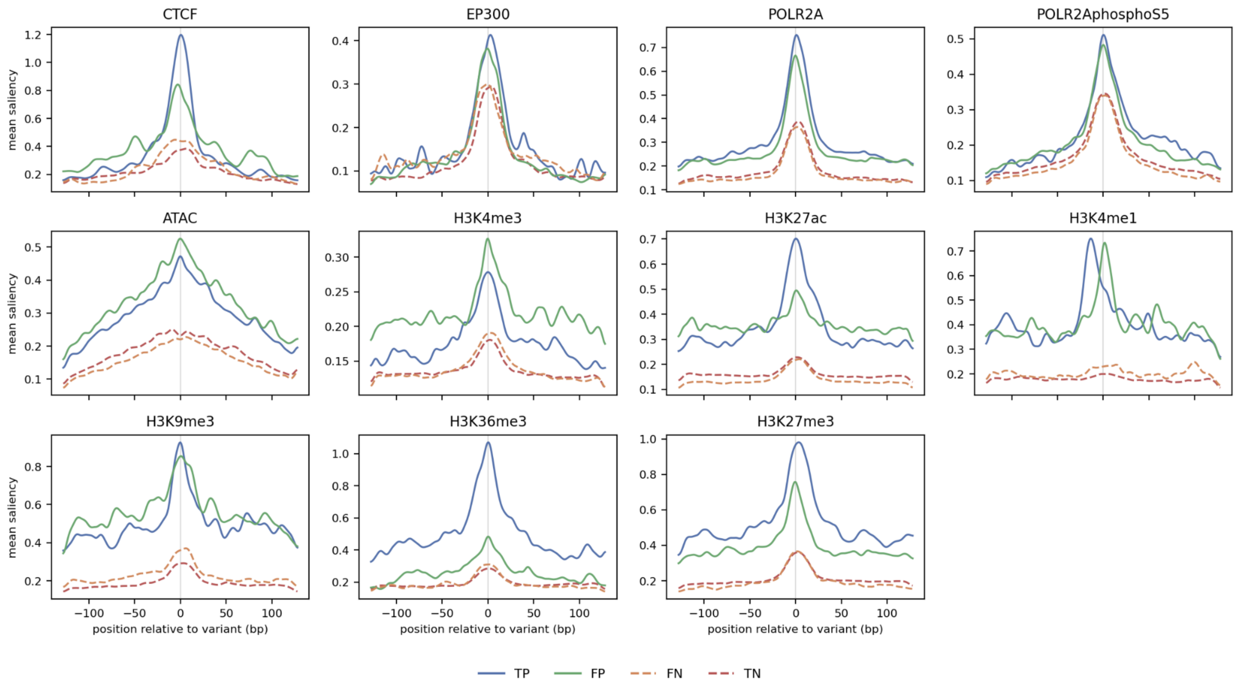


Figure S3. **ISM profiles averaged across confusion matrix outcomes from DNABERT-2 HapAdapter heads.** In silico saturation mutagenesis saliency profiles averaged over confusion matrix outcomes from the DNABERT-2 HapAdapter heads generally peak at the hetSNV of interest, with true positive classified sequences typically showing the highest central peak across assays and individuals. Displayed here just for individual 2 as we observe the same behavior for the other three individuals.
